# Heterogeneities in afferent connectivity dominate local heterogeneities in the emergence of response decorrelation in the dentate gyrus

**DOI:** 10.1101/173450

**Authors:** Poonam Mishra, Rishikesh Narayanan

## Abstract

The ability of a neuronal population to effectuate response decorrelation has been identified as an essential prelude to efficient neural encoding. To what extent are diverse forms of local and afferent heterogeneities essential in accomplishing such response decorrelation in the dentate gyrus (DG)? Here, we incrementally incorporated four distinct forms of biological heterogeneities into conductance-based network models of the DG and systematically delineate their relative contributions to response decorrelation. We incorporated intrinsic heterogeneities by stochastically generating several electrophysiologically-validated basket and granule cell models that exhibited significant parametric variability, and introduced synaptic heterogeneities through randomized local synaptic strengths. In including adult neurogenesis, we subjected the valid model populations to randomized structural plasticity and matched neuronal excitability to electrophysiological data. We assessed networks comprising different combinations of these three local heterogeneities with identical or heterogeneous afferent inputs from the entorhinal cortex. We found that the three forms of local heterogeneities were independently and synergistically capable of mediating significant response decorrelation when the network was driven by identical afferent inputs. Strikingly, however, when we incorporated afferent heterogeneities into the network to account for the unique divergence in DG afferent connectivity, the impact of all three forms of local heterogeneities were significantly suppressed by the dominant role of afferent heterogeneities in mediating response decorrelation. Our results unveil a unique convergence of cellular- and network-scale degeneracy in the emergence of response decorrelation in the DG, and constitute a significant departure from the literature that assigns a critical role for local network heterogeneities in input discriminability.

**SIGNIFICANCE STATEMENT:** The olfactory bulb and the dentate gyrus (DG) networks assimilate new neurons in adult rodents, with adult neurogenesis postulated to subserve efficacious information transfer by reducing correlations in neuronal responses to afferent inputs. Heterogeneities emerging from the lateral dendro-dendritic synapses, mediated by locally-projecting neurogenic inhibitory granule cells, are known to play critical roles in channel decorrelation in the olfactory bulb. However, the contributions of different heterogeneities in mediating response decorrelation in DG, comprising neurogenic excitatory granule cells projecting beyond DG and endowed with uniquely divergent afferent inputs, have not been delineated. Here, we quantitatively demonstrate the dominance of afferent heterogeneities, over multiple local heterogeneities, in the emergence of response decorrelation in DG, together unveiling cross-region degeneracy in accomplishing response decorrelation.

---

The ability of a neuronal population to effectuate response decorrelation has been identified as an essential prelude to efficient neural encoding, as the decorrelation process ensures that information conveyed by different neuronal channels is complementary (1-5). The critical importance of local circuit heterogeneities, — including those in intrinsic properties, in synaptic strengths and in neuronal structure, observed either under baseline conditions or achieved specifically through adult neurogenesis, — in achieving such response decorrelation has been recognized across different brain regions (1-18). Studies in the olfactory bulb (OB), one of the two prominent brain regions that express adult neurogenesis, have assessed the impact of these local heterogeneities on response decorrelation (1, 2, 6, 10), emphasizing the critical importance of intrinsic heterogeneities and lateral inhibition in the emergence of response decorrelation. However, despite the dentate gyrus (DG) being the other prominent brain region expressing adult neurogenesis and despite the widespread literature on the role of DG in pattern separation (7-9, 19-22), it is surprising that the impact of distinct forms of local and afferent heterogeneities on channel decorrelation has not been assessed in the DG.

This lacuna in the literature is especially striking because of the stark contrasts in terms of the unique afferent connectivity to the dentate and in the specific roles of adult neurogenesis in the DG *vs.* the OB (1-3, 7-9, 18-26), although both circuits have been implicated in response decorrelation and express adult neurogenesis. First, there is evidence for adult neurogenesis resulting in both *excitatory granule cells* and inhibitory basket cells in the dentate, whereas adult neurogenesis results in *inhibitory granule cells* in the OB. Second, OB granule cells lack axons and make *local lateral inhibitory dendrodendritic* synapses with other local circuit cells, and *do not* project outside the OB. In striking contrast, DG cells extend unmyelinated axons connecting *both within and beyond* (principally to CA3) the DG. It was this feature of the DG granule cells as the principal projection cell to the CA3 that was important in its theoretically postulated role in response decorrelation *before* inputs are fed to the pattern completing recurrent CA3 network (27, 28). Third, neurogenesis results in the replacement of the majority of granule neurons in the olfactory bulb, whereas it leads to a substantial addition of granule neurons to the hippocampal dentate gyrus, suggesting that adult neurogenesis could play distinct roles in the two brain regions (24). Finally, and most importantly, the principal inputs to the olfactory granule cells are the mitral cells through *local dendrodendritic synapses*, whereas the principal inputs to the ~1.2 billion dentate granule cells are the ~30,000 LII entorhinal cortical neurons. This significant divergence and sparsity of connections in the afferent projections from the excitatory cells in the EC to the excitatory DG granule cells is therefore unique to the DG, and is critical to the analysis in terms of the specific roles of local *vs.* afferent heterogeneities.

In the DG network, there are at least four distinct forms of heterogeneities that could mediate response decorrelation (the first three are local to the DG network whereas the fourth is afferent onto the network): (i) heterogeneity in intrinsic ion channel and excitability properties of the neurons; (ii) non-uniformities in the local synaptic connectivity; (iii) structural heterogeneities in neurons introduced by adult neurogenesis; and (iv) input-driven heterogeneity that is reflective of the distinct sets of afferent inputs that impinge on different neurons (as a consequence of the unique divergence in DG connectivity). Which of these distinct forms of heterogeneities play a critical role in mediating response decorrelation in the DG? Does a highly divergent, sparsely active network need an additional layer of neurogenesis-induced heterogeneity for effectuating response decorrelation? What is the impact of cell-to-cell variability in ion channel properties and excitability on response decorrelation in the DG network receiving different patterns of inputs?

In this study, we systematically and incrementally incorporate the four different forms of heterogeneities into conductance-based network models of the DG and delineate the impact of each form of heterogeneity on response decorrelation. Specifically, we employed a stochastic search algorithm spanning an exhaustive parametric space (involving experimentally-determined ion channel as well as neurophysiological properties) to reveal cellular-scale degeneracy in the DG, whereby disparate combinations of passive and active properties yielded analogous cellular physiology of excitatory granule (GC) and inhibitory basket cell (BC) populations. Next, we further expanded the parametric search space to encompass biologically observed heterogeneities in local/afferent network connectivity and in neurogenesis-induced alteration to neuronal structure and excitability. We systematically assessed response decorrelation in different DG networks, each built with incremental addition of the four distinct forms of heterogeneities. We found that in the absence of afferent heterogeneities, that is when the DG network was driven by identical afferent inputs, the three forms of local heterogeneities were independently and synergistically capable of mediating significant response decorrelation. Strikingly, however, when we incorporated afferent heterogeneities into the network to account for the unique divergence in DG afferent connectivity, we found that the impact of all three forms of local heterogeneities were suppressed by the dominant role played by afferent heterogeneities in mediating the emergence of response decorrelation. These conclusions on the dominance of afferent heterogeneities constitute a significant departure from the literature that assigns a critical role for local network heterogeneities (including those induced by adult neurogenesis) in input discriminability, and unveils crucial distinctions in the emergence of response decorrelation in the DG *vs.* the OB network.

## RESULTS

In systematically delineating the impact of distinct forms of heterogeneities on response decorrelation, we constructed networks of 500 GCs and 75 BCs from respective conductance-based model populations (Fig. *1A*–*B*). The heterogeneous conductance-based model populations of GC and BC neurons were derived from independent stochastic search procedures that replicated 9 different electrophysiological measurements (Fig. 1C–G) for each cell type (Tables S1–S4). These 575 cells were distributed in a cylindrical neuropil of 156-μm diameter and 40-μm depth (Fig. 1*A*), with cell density and local connection probability between GCs and BCs (Fig. 1*B*) matched with experimental equivalents. Each cell in the network received local circuit inputs from other BCs or GCs (Fig. 1*B*) and external inputs (Fig. 1*H*) from several cells in the medial (MEC) and lateral entorhinal cortices (LEC), which allowed it to fire (Fig. 1*I*) at specific locations (Fig. 1*J*) within the arena that the virtual animal traversed in randomized order (over the entire simulation period of 1000 s).

**Figure 1.**
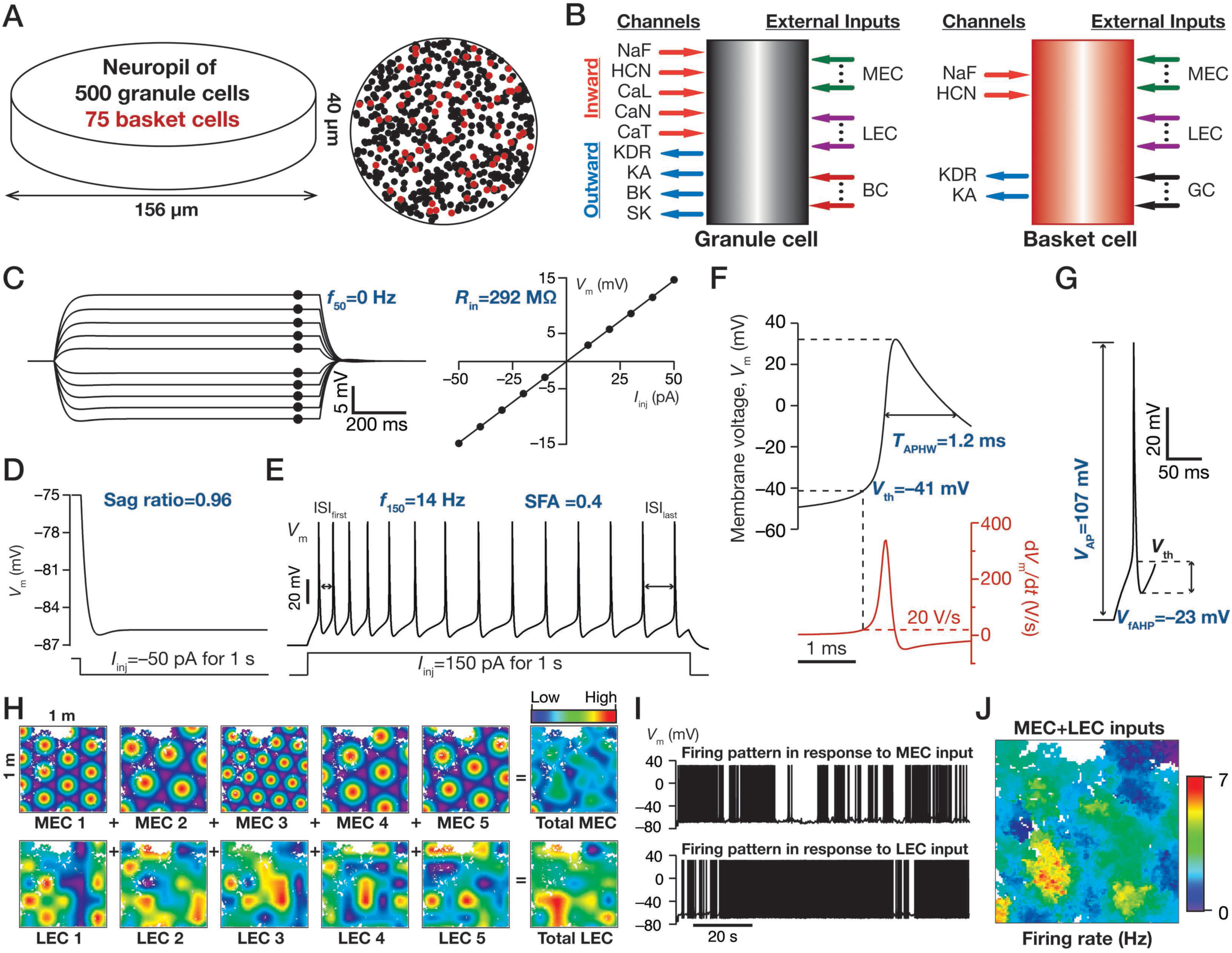
Model components and measurements. (*A*) Schematic representation of the cylindrical neuropil of 156 μm diameter and 40 μm height (left) with the top view (right) showing the distribution of 500 GCs (black) and 75 BCs (red). (B) Conductance-based models of GCs (left) and BCs (right) expressed different sets of ion channels and received external inputs from several MEC and LEC cells. (*C*–*G*) The nine physiological measurements employed in defining the GC populations: input resistance, *R*_in_, measured as the slope of a *V*–*I* curve obtained by plotting steady-state voltage responses to current pulses of amplitude –50 to 50 pA, in steps of 10 pA, for 500 ms (*C*); sag ratio, measured as the ratio between the steady-state voltage response and the peak voltage response to a –50 pA current pulse for 1 s (*D*); firing rate in response to 50 pA, *f*_50_ (*C*) and 150 pA current injection, *f*_150_ (*E*); spike frequency adaptation (SFA) computed as the ratio between the first (*ISI*_first_) and the last (*ISI*_las_t) inter-spike intervals in spiking response to a 150 pA current injection (*E*); action potential half-width, *T*_APHW_ (*F*); action potential threshold, computed as the voltage at the time point where d*V*_m_/d*t* crosses 20 V/s (*F*); action potential amplitude, *V*_AP_ (*G*) and the fast afterhyperpolarization potential (*V*_AHP_). (*H*) Inputs from MEC (top) were modeled as grid structures with randomized scale and orientation, whereas inputs from LEC (bottom), carrying contextual information, were represented as smoothed and randomized matrices comprised of active and inactive boxes. Schematic color-coded representations of individual inputs (5 MEC and 5 LEC cells) and their summations (separate for MEC and LEC inputs) are superimposed on the virtual animal trajectory in an arena of size 1 m × 1 m. (*I*) Sample GC voltage trace in response to total MEC (top) and LEC (bottom) current inputs. (*J*) Color-coded rate map obtained by superimposing firing rate output from an isolated GC in response to both MEC and LEC inputs, as the virtual animal traverses the arena.

### Degeneracy in single neuron physiology of granule and basket cell model populations

We employed a well-established stochastic search strategy (29-32) to arrive at populations of conductance-based models for GCs and BCs. This exhaustive parametric search procedure was performed on 40 parameters for GCs (Table S1), and 18 parameters for BCs (Table S3), involving ion channel properties derived from respective neuronal subtypes. Nine different measurements, defining excitability and action potential firing patterns (Fig. 1; Table S2), were obtained from each of the 20,000 stochastically generated unique GC models, and were matched with corresponding electrophysiological GC measurements. We found 126 of the 20,000 models (~0.63%) where all nine measurements were within these electrophysiological bounds (Table S2), and thus were declared as valid GC models. A similar procedure was employed for BC cells, where 9 different measurements from 8,000 unique models were compared with corresponding electrophysiological BC measurements. Here, we found 54 of the 8,000 models (~0.675%) where all nine measurements were within electrophysiological bounds (Table S4), and declared them as valid BC models. The experimental bounds on physiological measurements for granule (Table S2) and basket (Table S4) cells were obtained from references (33-37).

Did the validation process place tight restrictions on model parameters that resulted in the collapse of all valid models to be near-homogeneous equivalents with very little changes in their parametric values? To address this, we plotted model parameters of 6 GCs (Fig. 2) and 6 BCs (Fig. S1), which had near-identical measurements values, and found the parametric values to spread through a wide span of the range employed in the respective stochastic searches. To further validate this, we plotted histograms of each of the 40 GC model parameters and the 18 BC model parameters, and found them to spread through the entire span of their respective ranges (Fig. 3*A*). These results demonstrated that the valid models were not near-homogeneous parametric equivalents, but form heterogeneous populations of GCs and BCs that functionally matched their respective electrophysiological measurements, thereby unveiling cellular-scale degeneracy in GC and BC neurons.

**Figure 2.**
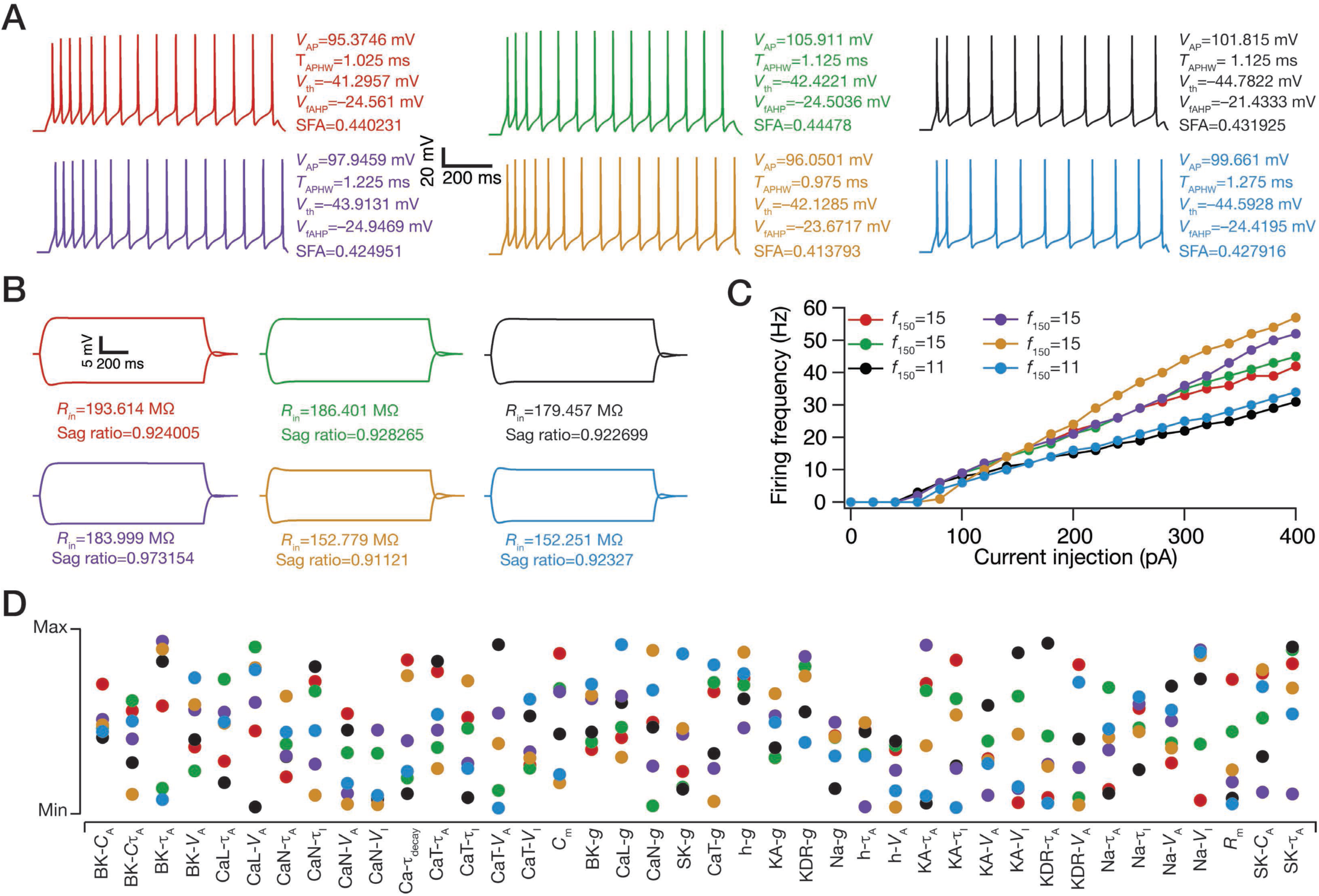
Illustration of cellular-scale degeneracy in granule cell physiology with six randomly chosen valid models, where analogous functional characteristics were achieved through disparate parametric combinations. (*A*) Firing pattern of six randomly chosen valid GC models in response to 150 pA current injection with corresponding measurement values for action potential amplitude (*V*_AP_), action potential half-width (*T*_APHW_), action potential threshold (*V*_th_), fast afterhyperpolarization (*V*_fAHP_) and spike frequency adaptation (SFA). (*B*) Voltage traces of six valid GC models in response to –50 pA and 50 pA current injection, with associated measurement values for input resistance (*R*_in_) and sag ratio. Note that firing rate at 150 pA, *f*_50_, was zero for all models. (*C*) Firing frequency plots for six valid GC models in response to 0–400 pA current injections, indicating values of firing rate at 150 pA for each valid model. Note that all the 9 different measurements are very similar across these six models. (*D*) Distribution of the 40 underlying parameters in the six valid models, shown with reference to their respective min–max ranges. The color code of the dots is matched with the plots and traces for the corresponding valid models in (*A*)–(*C*).

**Figure 3.**
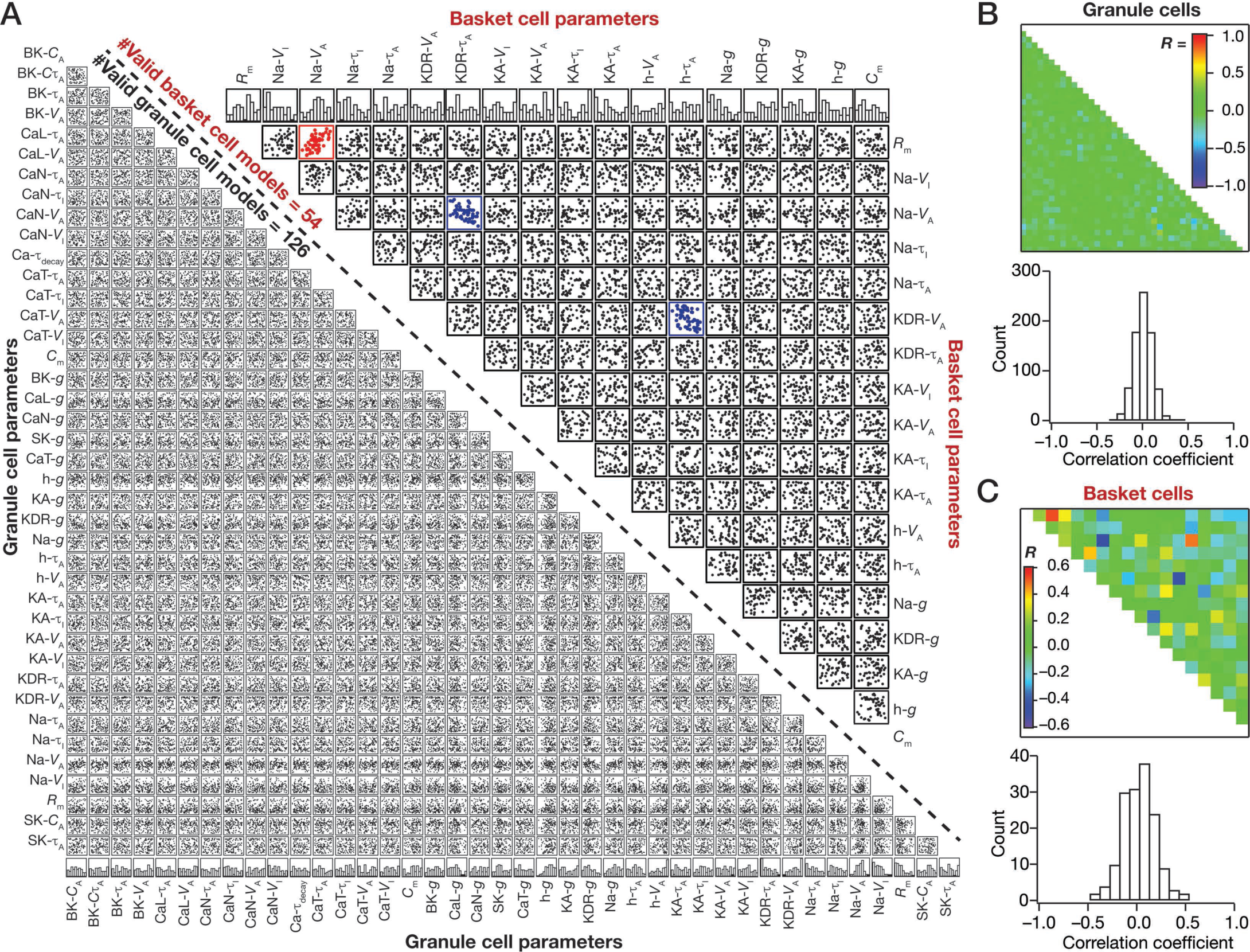
Independently heterogeneous populations of granule and basket cells exhibited cellular-scale degeneracy with weak pair-wise correlations of underlying parameters. (*A*) *Left*, Lower triangular part of a matrix comprising pair-wise scatter plots between 40 parameters underlying all valid GC models (*n*=126). The bottommost row represents the histograms for corresponding parameters in the valid model population, showing all parameters spanning their respective min-max ranges. *Right*, Upper triangular part of a matrix comprising pair-wise scatter plots between 18 parameters underlying all valid BC models (*n*=54). The topmost row represents the histograms for corresponding parameters in the valid model population, showing all parameters spanning their respective min-max ranges. The red scatter plots indicate that the value of correlation coefficient for the pair was greater than 0.5, whereas the blue scatter plots denote pairs where the correlation coefficient value was less than –0.5. (*B*) *Top*, Heat map of correlation coefficient values for GC cells, corresponding to each scatter plot box depicted in (*A*). *Bottom*, Distribution of correlation coefficient values for the 780 unique pairs, of the 40 parameters, corresponding to scatter plots for GC parameters shown in (*A*). (*C*) Same as (*B*) but for BC cells with 153 unique pairs of correlation coefficients (*A*).

How did these neuronal populations achieve degeneracy? Did they achieve this by pair-wise compensation across parameters, or was change in one parameter compensated by changes in several other parameters to achieve robust physiological equivalence? In answering this, we plotted pair-wise scatter plots, independently on valid model parameters of the GC and BC populations (Fig. 3*A*), and computed pair-wise Pearson’s correlation coefficients for each scatter plot (Fig. 3*B*–*C*). We found that a vast majority of these pairs displayed very weak pair-wise correlations (*R*^2^ < 0.25; Fig. 3*B*–*C*), suggesting that degeneracy in both populations was achieved through collective changes spanning several parameters.

### Heterogeneities in neuronal intrinsic properties mediated decorrelation of neuronal responses to *identical* external inputs

Cellular-scale degeneracy in these valid model populations provided an ideal manifestation of physiologically constrained intrinsic heterogeneities in the GC and BC model populations. Consequently, in defining the first layer of heterogeneity, we constructed a network of these heterogeneous populations with *identical* external inputs from the MEC and LEC and homogenous local synaptic connectivity.

We allowed the virtual animal to traverse the arena, recorded the voltage traces of all the GCs and BCs in this network, computed their firing rates and overlaid neuronal firing structure on the arena to observe the emergence of place fields (Fig. 4*A*). To quantify the extent of decorrelation achieved through the introduction of intrinsic heterogeneities, we computed instantaneous firing rates of all neurons in the network across the entire traversal period (Fig. 4*A*; Fig. S2) and calculated pair-wise Pearson’s correlation coefficients across these firing rate arrays for all neurons (Fig. 4*B*; Fig. S2B). If the network were composed of a homogeneous population of GCs and BCs receiving *identical* inputs, then the responses of all GCs would be identical to each other, with all pair-wise correlation coefficients set at unity. However, owing to heterogeneous intrinsic excitability of individual neurons, their responses exhibited significant differences, especially in terms of overall firing rate at individual place fields (Fig. 4*A*; Fig. S2), even with *identical* external inputs and homogeneous local synaptic weights. Such dissimilarity in neuronal firing rate response emerges from two distinct manifestations of intrinsic heterogeneity. First, certain periods of identical synaptic inputs would be subthreshold for neurons with lower excitability (*e.g.*, Cell #2 in Fig. 4*A*), but would be suprathreshold for neurons with relatively higher excitability (*e.g.*, Cell #5 in Fig. 4*A*), thereby manifesting as changes in firing rate or in the emergence of place fields at specific locations (38). Additionally, these observations also suggest that DG neurons could undergo rate remapping (22, 26) merely as a consequence of plasticity in intrinsic excitability. Second, although the numbers and synaptic weights of excitatory or inhibitory synapses received by neurons were identical, the patterns of activation of these synapses would be different across neurons as a consequence of significant variability in their respective presynaptic neuronal firing (Fig. 4*A*, Fig. S2).

**Figure 4.**
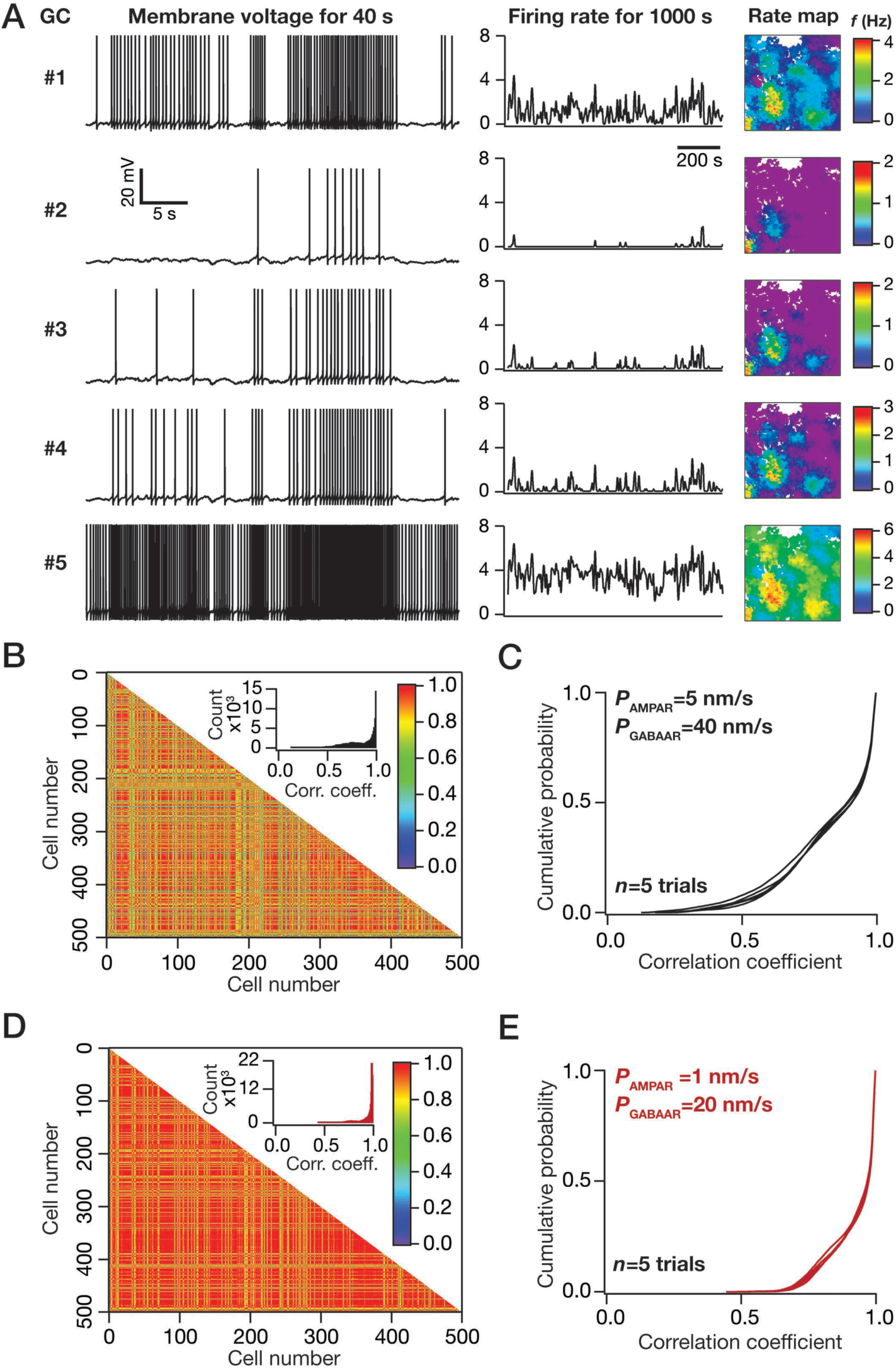
Heterogeneity in intrinsic neuronal excitability is a robust mechanism for achieving response decorrelation through rate remapping of cellular responses. (*A*) Voltage traces (left), instantaneous firing rate (middle) and color-coded rate maps (right; superimposed on the arena) for five different GCs in a network made of a heterogeneous GC and BC populations. (*B*) Lower triangular part of correlation matrix representing pair-wise Pearson’s correlation coefficient computed for firing rates of 500 GCs spanning the entire 1000 s simulation period. Inset represents the histogram of these correlation coefficients. Note that there was no heterogeneity in the synaptic strengths of local connections, with *P*_AMPAR_=5 nm/s and *P*_GABAAR_=40 nm/s for all excitatory and inhibitory synapses, respectively. (*C*) Cumulative distribution of correlation coefficients represented in matrix in (*B*). Plotted are distributions from five different trials of the simulation, with each trial different in terms of the cells picked to construct the network. (*D*–*E*) Same as (*B*–*C*), but with the synaptic strengths of local connections fixed at lower permeability values: *P*_AMPAR_=1 nm/s and *P*_GABAAR_=20 nm/s.

Consequent to such variability in firing responses of this intrinsically heterogeneous population of neurons, we found the distribution of correlation coefficients of instantaneous firing rates to be significantly (Kolmogorov Smirnov, KS test; *p*<0.001) different from an all-unity distribution representative of identical responses achieved in the absence of intrinsic variability (Fig. 4*B*–C; Fig. S2C). Next, we repeated these simulations with different combinations of excitatory and inhibitory synaptic weights, setting all local synapses to the same value, and computed cumulative histograms of firing rate correlation coefficients (Fig. 4*D*–*E*; Fig. S2*D*–*E*). We found a significant shift (Fig. 4*C* *vs.* Fig. 4*E*; KS test; *p*<0.001) in the level of decorrelation with different combinations of synaptic weights.

### Synaptic heterogeneity modulates decorrelation of neuronal responses to *identical* external inputs

Motivated by observations on the role of the local synaptic weights in modulating response decorrelation, we systematically assessed the impact of altering the excitatory and inhibitory synaptic weights on the correlation histograms. As a first step, the network was endowed with intrinsic heterogeneities and all local synaptic weights were identical but were assigned different values across different simulations (Fig. 5*A*–*B*; Fig. S3*A*–*B*). Although increases in either excitatory or inhibitory weights significantly enhanced the level of response decorrelation, the impact of increasing inhibitory weights had a dominant impact on decorrelating network responses (Fig. 5*A*–*B*; Fig. S3*A*–*B*) emphasizing the critical role of local inhibitory neurons in defining response decorrelation in excitatory neurons (9, 12, 13).

**Figure 5.**
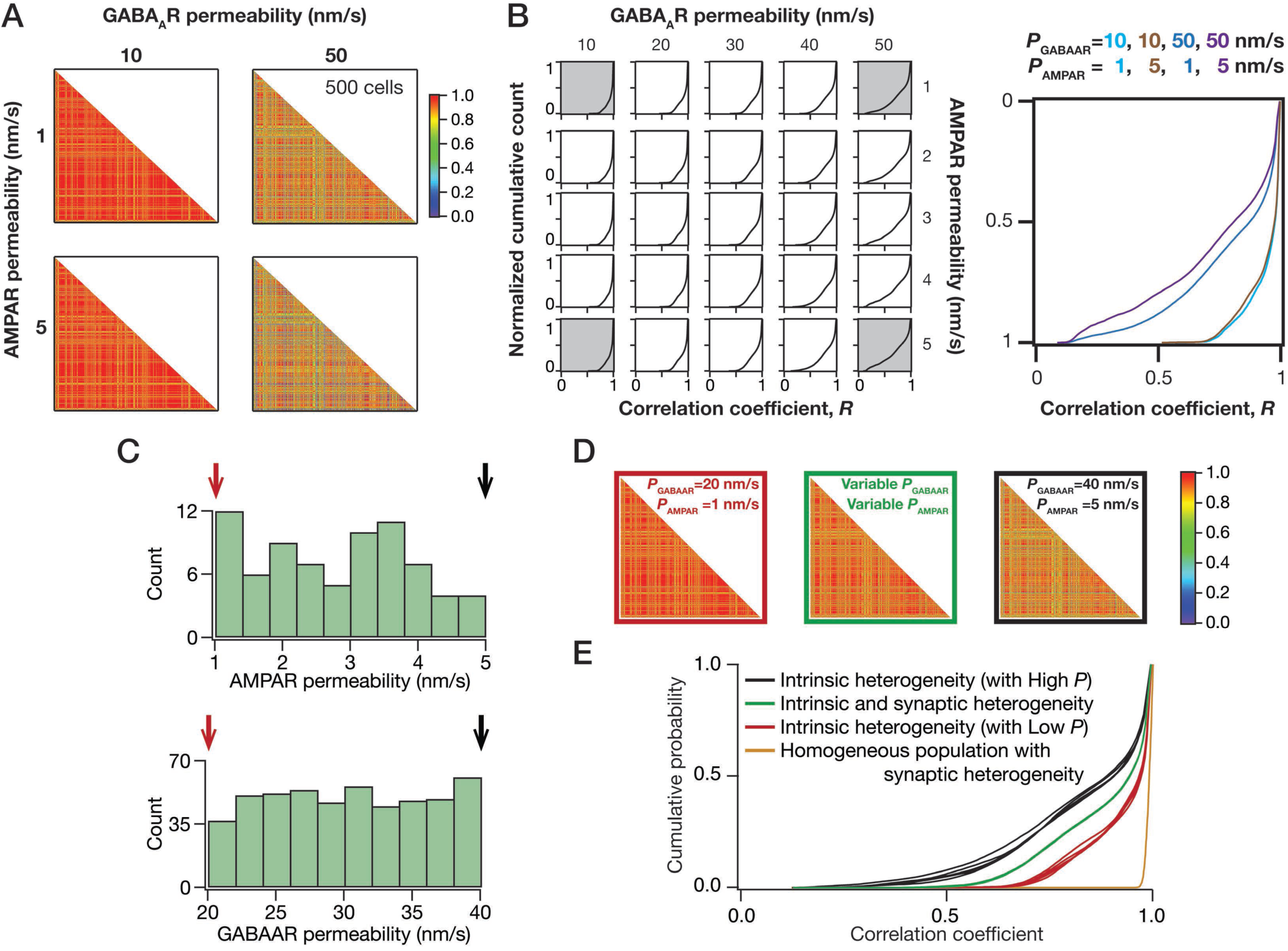
Heterogeneities in the strength of local network connections modulate response decorrelation, with increase in inhibitory synaptic strength enhancing network decorrelation. (*A*) Lower triangular part of correlation matrix representing pair-wise Pearson’s correlation coefficient computed for firing rates of 500 GCs. Note that there was no heterogeneity in the synaptic strengths of local connections, with AMPAR and GABAAR permeability across local network synapses set at fixed values. Shown are four different correlation matrices, with *P*_AMPAR_ (1 or 5 nm/s) and *P*_GABAAR_ (10 or 50 nm/s) fixed at one of the two values. (*B*) *Left*, Cumulative distribution of correlation coefficients for firing rates of 500 GCs, computed when the simulations were performed with different sets of fixed values of *P*_AMPAR_ (spanning 1–5 nm/s) and *P*_GABAAR_ (spanning 10–50 nm/s). The gray-shaded plots on the extremes were computed from corresponding matrices shown in (*A*). *Right*, Cumulative distributions of correlation coefficients corresponding to the gray-shaded plots on the left, to emphasize the impact of synaptic heterogeneity on decorrelation. (*C*) Distribution of *P*_AMPAR_ and *P*_GABAAR_ in a network of heterogeneous GC and BC populations, constructed with heterogeneity in local synaptic strengths as well. Each AMPA and GABA_A_ receptor permeability was picked from a uniform distribution that spanned the respective ranges. The color codes of arrows and plots correspond to cases plotted in (*D*)–(*E*). (*D*) Lower triangular part of correlation matrices representing pair-wise Pearson’s correlation coefficient computed for firing rates of 500 GCs. For the left and right matrices, which are the same plots as in Fig. 4*E* and Fig. 4*C*, respectively, there was no synaptic heterogeneity, with P_AMPAR_ and *P*_GABAAR_ set at specified fixed values for all excitatory and inhibitory synapses. The matrix represented in the center was computed from a network endowed with intrinsic as well as synaptic heterogeneity (shown in *C*). (*E*) Cumulative distribution of correlation coefficients represented in matrices in (*E*). Plotted are distributions from five different trials of each configuration. Note that except for the homogenous population, all three configurations were endowed with intrinsic heterogeneity. The configurations “Intrinsic + synaptic heterogeneity” and “Homogeneous + synaptic heterogeneity” had randomized synaptic permeabilities; for the other two configurations, the synaptic strengths were fixed at specific values: High P, *P*_AMPAR_=5 nm/s and *P*_GABAAR_=40 nm/s; Low P, *P*_AMPAR_=1 nm/s and *P*_GABAAR_=20 nm/s.

Would introduction of synaptic heterogeneities, where different synapses in the local network assume distinct values, further enhance neuronal response decorrelation? To test this, we assigned weights of excitatory and inhibitory synapses in the local network to randomized values picked from respective uniform distributions (Fig. 5*C*–*E*; Fig. S3*C*). Surprisingly, we found that introduction of synaptic heterogeneity did not enhance the level of response decorrelation, but allowed response decorrelation to express at a level that was within the bounds set by extreme values of identical synaptic weights (Fig. 5*E*; Fig. S3*C*). Importantly, the level of decorrelation achieved by the introduction of local synaptic heterogeneity into a homogeneous population (no intrinsic heterogeneity) of GCs and BCs was significantly minimal compared to that achieved by the mere presence of intrinsic heterogeneity (Fig. 5*E*; Fig. S3C; *cf.* Fig. 4, Fig. S2). Together, although the introduction of synaptic heterogeneity critically modulated the level of response decorrelation, these results suggest intrinsic heterogeneity as the dominant form among intrinsic and synaptic forms of heterogeneities in mediating response decorrelation.

### Adult neurogenesis-induced structural heterogeneity in neuronal age enhances decorrelation of neuronal responses to *identical* external inputs

A prominent hypothesis on the specific functions of adult neurogenesis in DG neurons is on their role in response decorrelation. One part of the rationale behind this hypothesis is the distinct excitability properties of new neurons that provide an additional layer of heterogeneity (7-9, 15, 19-21, 39). Although there are lines of evidence linking adult neurogenesis to response decorrelation, the specific role of new neurons and the additional layer of heterogeneity introduced by them in regulating input discriminability has not been systematically assessed.

To introduce neurogenesis-induced heterogeneity into our network, we noted that the excitability of new born neurons in the DG, which could mature to either GCs or BCs, is higher as a consequence of lower surface area reflective of the diminished arborization of immature neurons (9, 15, 39, 40). To quantitatively match the excitability properties of these neurons, we introduced structural plasticity by reducing the surface area of the valid GC and BC models (Fig. 3) through reduction of their diameter. This reduction in surface area expresses as an increase in input resistance (39, 41, 42) in each of these neurons (Fig. 6*A*), which in turn translates to increase in firing rate (Fig. 6*B*).

**Figure 6.**
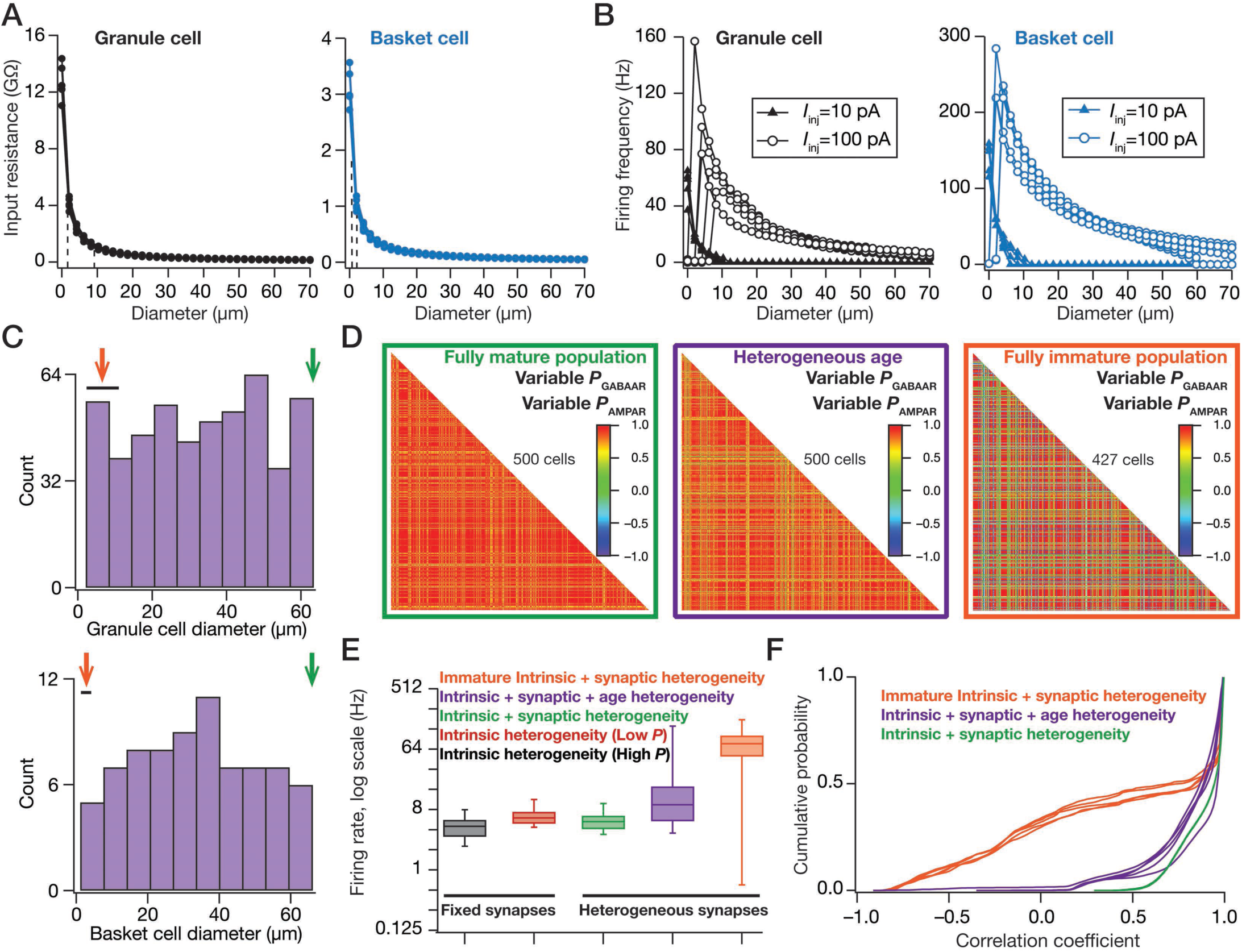
Incorporation of neurogenesis-induced structural heterogeneity in neuronal age enhances response decorrelation in a network of neurons receiving *identical* inputs. (*A*) Input resistance of the 126 GCs (left) and 54 BCs (right) plotted as a function of diameter of cell. Dotted lines represent the range for immature cell diameters (2–9 μm for GC and 1–3 μm for BC), obtained from ranges of experimentally obtained input resistance values in immature cells. (*B*) Firing frequency plotted as a function of diameter in response to 10 pA (closed triangles) and 100 pA (open circles) current injections into the 126 GCs (left) and 54 BCs (right). (*C*) Distribution of GC (top) and BC (bottom) diameters in a network of heterogeneous GC and BC populations, constructed with heterogeneity in local synaptic strengths and in the age of the neurons. The diameter of each GC and BC in the network was picked from a uniform distribution that spanned respective ranges. The color codes of arrows and plots correspond to fully mature network (green; large diameters), fully immature network (orange; small diameters) and mixed network (purple; variable diameters) cases plotted in (*D*)–(*F*). (*D*) Lower triangular part of correlation matrices representing pair-wise Pearson’s correlation coefficient computed for firing rates of all GCs. The matrix corresponding to the fully mature population is the same as that in Fig. 5*D*, with the same color code. Note that all three networks were endowed with intrinsic as well as synaptic heterogeneity, with changes only in the neuronal age. (*E*) Firing rates, represented as quartiles, of all GCs plotted for the different networks they resided in. (*F*) Cumulative distribution of correlation coefficients represented in matrices in (*D*). Plotted are distributions from five different trials of each configuration.

With the ability to introduce intrinsic, synaptic and neurogenesis-induced forms of heterogeneity into our network, we analyzed three distinct networks (fully mature, fully immature and variable age) to specifically understand the role of neurogenesis-induced heterogeneity on response decorrelation to *identical* inputs. All three networks were endowed with intrinsic as well as synaptic heterogeneities receiving afferent inputs from the same arena (Figs. 4–5), and the distinction between the three cases was only with reference to neuronal age (Fig. 6*D*). In comparing the firing rates of the GCs for different network configurations, we found that the firing rates of all GCs were comparable for all cases where neurogenesis-induced heterogeneities were absent. However, with the introduction of neurogenesis, especially in the scenario where all cells were immature, the firing rates increased and spanned a larger range. In the more physiologically relevant scenario of heterogeneous cellular age, although the firing rates spanned a larger range, a significant proportion of them were in the low firing regime characteristic of GCs (Fig. 6*E*).

We found that the level of response decorrelation in the fully immature network was significantly (KS test; *p*<0.001) higher than that achieved in the fully mature network (Fig. 6*F*). This is to be expected because the structural heterogeneity (effectuated by changes in diameter) would amplify the inherent intrinsic heterogeneity of neurons in the network, thereby enhancing the beneficiary effects of intrinsic heterogeneity that we had observed earlier (Fig. 4). Importantly, reminiscent of our results with the introduction of synaptic heterogeneity (Fig. 5), in the network that was endowed with variability in neuronal age, the level of decorrelation was intermediate between that obtained with the fully mature and the fully immature networks (Fig. 6*F*). Together, these results demonstrate that neurogenesis-induced variability in neuronal response properties adds an additional layer of heterogeneity in the DG network, and enhances network decorrelation to *identical* external inputs.

### Input-driven heterogeneity mediated by sparseness of afferent connectivity is a dominant regulator of neuronal response decorrelation

An important defining characteristic of the DG network is the sparseness of the afferent connectivity matrix that is reflective of massive convergence and divergence reflecting the small number of layer II EC cells (~30,000) that project to a large (~1.2 million) number of DG cells, resulting in significant variability in the set of afferent external inputs impinging on each GC (23, 43). Thus far in our analysis, in an effort to delineate the impact of three distinct forms of heterogeneity, we employed an artificial construct where all neurons in the network received *identical* inputs. To assess the impact of this fourth form of afferent input-driven heterogeneity, we introduced divergence in the set of EC neurons that project onto each GC and BC. This implied that each GC and BC now received distinct sets of EC inputs.

As a consequence of distinct set of inputs impinging on each GC, the firing fields were distinct across different GCs (Fig. 7*A*) and BCs (Fig. S4), unlike the near-identical firing fields (except for differences in firing frequency or threshold) in the case where neurons received *identical* inputs (Fig. 4*A*; Fig. S2). Importantly, when we analyzed pair-wise correlation of firing rates across different neurons, we found that the correlation coefficients were lower irrespective of the presence or absence of different forms of heterogeneity (Fig. 7*B*). The overall firing rate distributions obtained with either *identical* (Fig. 6) or *distinct* (Fig. 7*C*) afferent inputs were similar, thereby ruling out changes in firing rate as a possible cause for the observed differences in correlation coefficients.

**Figure 7.**
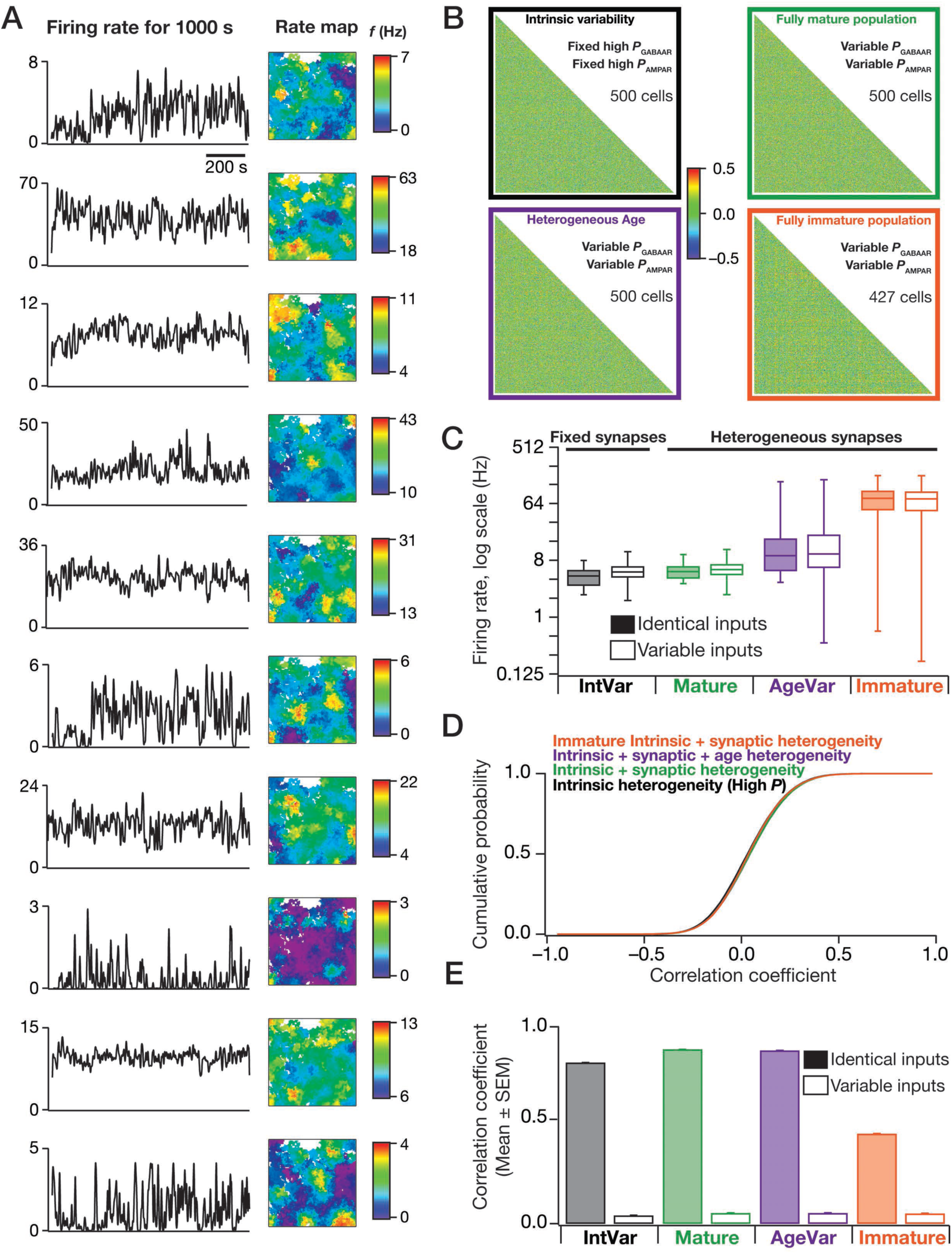
Heterogeneous external connectivity is the dominant form of variability that drives response decorrelation in a network endowed with intrinsic, synaptic and age heterogeneities. (*A*) Instantaneous firing rate (left) and color-coded rate maps (right; superimposed on the arena) for ten different GCs in a network endowed with intrinsic, synaptic, age and input-driven forms of heterogeneities. (*B*) Lower triangular part of correlation matrices representing pair-wise Pearson’s correlation coefficient computed for firing rates of all GCs. The four different matrices correspond to networks endowed with different sets of heterogeneities. (*C*) Firing rates, represented as quartiles, of all the GCs plotted for the different networks they resided in. Color-codes for the specific set of heterogeneities included into the network are the same as those in panel *B* above. (*D*) Cumulative distribution of correlation coefficients represented in matrices in (*B*). (*E*) Statistical (mean ± SEM) comparison of correlation coefficients obtained with networks, endowed with distinct forms of heterogeneities, receiving *identical* (solid boxes; derived from Fig. 6*F*) *vs.* variable (open boxes; derived from panel *D*) external inputs.

Strikingly, when we plotted the cumulative distributions of correlation coefficients obtained with the introduction of distinct forms of network heterogeneities, we found them to significantly overlap with each other (Fig. 7*D*). This is in stark contrast to the network receiving *identical* external inputs (Fig. 5*E*, Fig. 6*F*), where introduction of each of intrinsic, synaptic and neurogenesis-induced heterogeneities enhanced or altered the level of response decorrelation. The negligible impact of the intrinsic or synaptic or age heterogeneities on the overall level of response decorrelation achieved in the presence of input-driven heterogeneities, which was higher than that obtained with *identical* inputs (Fig. 7*E*), unveiled the dominance of heterogeneities driven by afferent connectivity in determining response decorrelation.

Were our conclusions on the role of different forms of heterogeneities scalable and invariant to network size? To test this, we repeated our analyses in Figs. 4–7 with a smaller network made of 100 GCs and 15 BCs, and found our conclusions to scale across different network sizes (Fig. S5). Together, our results demonstrate that local heterogeneities in intrinsic, synaptic and neuronal structural (driven by adult neurogenesis) properties contributed to significant levels of response decorrelation in the presence of *identical* afferent synaptic drive. However, when the network received heterogeneous external inputs, the impact of local heterogeneities on response decorrelation was strongly suppressed by the dominant role of afferent heterogeneities in mediating neuronal response decorrelation.

## DISCUSSION

### Dominance of input-driven heterogeneity and implications for the physiological roles for adult neurogenesis

Our results quantitatively demonstrate a dominant role for afferent heterogeneities, driven specifically by the unique network structure of the DG involving huge number of GCs innervated by the small number of LII EC neurons, in driving response decorrelation in the DG. Within this framework, this dominant connectivity divergence in feed-forward afferents, synergistically coupled to the hetereogeneous intrinsic properties and the sparse GC activity that is sharpened by the local inhibitory network places the DG network as an ideal decorrelating system. Importantly, our conclusions on the dominance of heterogeneous afferent connectivity, with the local network heterogeneities playing secondary roles, pose specific questions on the role of adult neurogenesis in input discriminability, questioning the rationale behind the need for new neurons in a highly divergent, sparsely active network (9).

If adult neurogenesis-induced heterogeneities in neuronal properties were not the dominant contributor to response decorrelation, what is the precise role of adult neurogenesis in the DG? One possibility within our framework is that adult neurogenesis could be a mechanism for implementing afferent heterogeneities across DG neurons, whereby new neurons establish connections to afferent fibers in an activity-dependent manner (14, 44-46), thereby assigning a specific set of active afferent inputs to new neurons of the same time of birth (9, 25, 47). In such a scenario, the afferent heterogeneities would be driven by active assignment of spatial connectivity from the EC to individual DG neurons, whereby the novel contexts encountered by the animal are encoded by the temporal onset of neurons. Such active assignments could be driven by activity-dependent connectivity aided by the hyper-plastic, hyper-excitable nature of new neurons, and the resultant afferent heterogeneities (different neurons get different EC inputs) then plays specific roles in response decorrelation, in encoding temporal context and in controlling memory resolution (9, 14, 20, 25, 39, 44-47). In addition to this, our results suggest that the variability introduced by new neurons in terms of their intrinsic excitability (Figs. 4, 6) and in terms of altered excitation-inhibition balance (Fig. 5) could also be candidate mechanisms through which adult neurogenesis enhances (beyond what is driven by afferent heterogeneities) the degree of response decorrelation achieved in the DG network (8, 9, 16, 20, 25, 47).

### Multiscale degeneracy: Convergence of different scales of degeneracy to achieve single-neuron homeostasis and response decorrelation

A central premise of robustness in biological function is degeneracy, where distinct structural components could combine to elicit analogous function. Given the several possible routes through which similar function can be achieved, it is possible for biological systems to invoke compensatory mechanisms to achieve the same function through very different parametric combinations (29-32, 48). Complementary to this powerful theory, our results show that identical levels of output *dissimilarity* could be achieved with disparate classes of parametric heterogeneity. Specifically, whereas degeneracy suggests that *similar* functionalities could be achieved with disparate combinations of constituent parameters, our results argue for a case where similar degrees of response decorrelation could be achieved through disparate classes of heterogeneity. Do our results constitute a departure from the premise of degeneracy in biological systems?

In systems that are responsible for encoding of novel information, robust homeostasis of output constitutes only one side of the overall physiological goals. The other side constitutes encoding of new information, which by definition involves changes to certain output characteristics to reflect this encoding process. With specific reference to the DG, whose prime *encoding* function has been postulated to be response decorrelation, it is important that the focus is not on mere maintenance of robust outputs. If response decorrelation were the overall function, and different classes of heterogeneity are considered as disparate structural components, our conclusions are consistent with the overall framework of degeneracy where distinct structural components could come together to elicit analogous function. Thus, there are several layers of degeneracy, spanning the molecular, cellular, network and behavioral scales of analyses, embedded in results presented in this study. At the cellular scale, distinct combinations of intrinsic parameters (the molecular components: the ion channels) come together to elicit analogous cellular response properties. At the network scale, distinct combinations of intrinsic and synaptic properties interact to elicit similar levels of response decorrelation. Together, our results unveil a systematic convergence of degeneracy spanning different scales of analysis in the DG network, achieving the twin goals of the DG network (input discriminability and firing rate homeostasis) within the broad framework of degeneracy.

### Comparison of mechanisms for decorrelation in the dentate gyrus and in the olfactory bulb

The olfactory bulb (OB) is another brain region that expresses adult neurogenesis and has been postulated to play a critical role in channel decorrelation (referred here as response decorrelation) (1-3, 18). Although there are similarities in our conclusions with those in the olfactory literature in terms of the roles of neuronal nonlinearites, intrinsic heterogeneities and inhibition in effectuating channel decorrelation in the *absence* of afferent heterogeneities, the significant departure in our conclusions is with reference to the dominant role of afferent heterogeneities. We argue the dominance of afferent heterogeneities is a distinctive feature of the DG circuit, and is reflective of the unique afferent connectivity to the dentate and the several stark contrasts between the roles of adult neurogenesis in the DG *vs.* the OB circuit (1-3, 7-9, 18-26). Specifically, the significant divergence and sparsity of connections in the afferent projections from the excitatory cells in the EC to the excitatory GCs is unique to the DG, and is critical to the conclusions that we draw here in terms of the dominance of the afferent heterogeneities in mediating response decorrelation.

The well-established distinctions between these two networks, in conjunction with our conclusions on the unique role of afferent heterogeneities in the DG network suggest that the mechanisms behind achieving decorrelation in the OB and the DG networks are very different. Specifically, whereas decorrelation in the OB has been postulated to be aided by new laterally inhibiting neurons forming dendrodendritic synapses across the local circuit (1-3, 18), our conclusions here, derived from the specifics of the DG network, its intrinsic heterogeneities and its afferent connectivity, present a dominant role for the afferent heterogeneities supported by synergistic interactions with local heterogeneities. Together, these disparate structural routes to achieve decorrelation further emphasize our conclusions on degeneracy in encoding mechanisms. In discussing the role of distinct forms of heterogeneities in effectuating decorrelation, we had emphasized in the previous section the possibility of how distinct forms of heterogeneities could be recruited to achieve analogous levels of decorrelation. With these distinctions between the OB and the DG, it is clear that this degree of degeneracy could be much broader where the OB and DG could be using adult neurogenesis in very different ways towards achieving response decorrelation.

## METHODS

Detailed experimental procedures are provided in the SI Methods. Briefly, the principal goal of this study was to systematically assess the impact of different forms of heterogeneities on response decorrelation in the DG. In doing this, we took advantage of the versatility of conductance-based neuronal network models, and distinguished between four different types of heterogeneities: (i) *intrinsic heterogeneity*, where the GC and BC model neurons that were employed to construct the network had widely variable intrinsic parametric combinations yielding physiological measurements that matched their experimental counterparts. These heterogeneous model populations were obtained using independent stochastic search procedures for GCs and BCs; (ii) *synaptic heterogeneity*, where the synaptic strength of the local GC-BC network was variable with excitatory and inhibitory synaptic permeability values picked from uniform random distributions; (iii) *neurogenesis*-*induced heterogeneity in age* of the neuron, where the DG network could be made entirely of mature or immature neurons, or be constructed from neurons that represented different randomized neuronal ages; and (iv) *input*-*driven heterogeneity*, where the GC and BC populations received either *identical* inputs from the EC, or each GC and BC received unique inputs from the EC. Networks endowed with different combinations of heterogeneities were constructed, and received inputs that were tied to the location of a virtual animal traversing in an arena. Spike timings of individual neurons in response to arena traversals were recorded, and pairwise correlations were computed on instantaneous firing rates of GC or BC neuron populations. Analyses of correlation coefficients across distinct networks endowed with disparate forms of heterogeneities were employed to derive insights into the roles of distinct forms of heterogeneities in driving input discriminability in these networks. All simulations were performed using the NEURON simulation environment (49).

## Acknowledgments

This work was supported by the Wellcome Trust-DBT India Alliance (Senior fellowship to RN; IA/S/16/2/502727), Human Frontier Science Program (HFSP) Organization (RN), the Department of Biotechnology (RN), the Department of Science and Technology (RN), and the Ministry of Human Resource Development (RN & PM). The authors thank the members of the cellular neurophysiology laboratory for helpful discussions and for comments on a draft of this manuscript.

## REFERENCES

1. Wiechert MT, Judkewitz B, Riecke H, & Friedrich RW (2010) Mechanisms of pattern decorrelation by recurrent neuronal circuits. Nat Neurosci 13(8):1003-1010.

2. Padmanabhan K & Urban NN (2010) Intrinsic biophysical diversity decorrelates neuronal firing while increasing information content. Nat Neurosci 13(10): 1276-1282.

3. Chow SF, Wick SD, & Riecke H (2012) Neurogenesis drives stimulus decorrelation in a model of the olfactory bulb. PLoS computational biology 8(3):e1002398.

4. Pitkow X & Meister M (2012) Decorrelation and efficient coding by retinal ganglion cells. Nat Neurosci 15(4):628-635.

5. Tetzlaff T, Helias M, Einevoll GT, & Diesmann M (2012) Decorrelation of neural-network activity by inhibitory feedback. PLoS computational biology 8(8):e1002596.

6. Luo SX, Axel R, & Abbott LF (2010) Generating sparse and selective third-order responses in the olfactory system of the fly. Proc Natl Acad Sci U S A 107(23): 10713-10718.

7. Aimone JB, Deng W, & Gage FH (2010) Adult neurogenesis: integrating theories and separating functions. Trends Cogn Sci 14(7):325-337.

8. Aimone JB, Deng W, & Gage FH (2011) Resolving new memories: a critical look at the dentate gyrus, adult neurogenesis, and pattern separation. Neuron 70(4):589-596.

9. Aimone JB, Li Y, Lee SW, Clemenson GD, Deng W, & Gage FH (2014) Regulation and function of adult neurogenesis: from genes to cognition. Physiol Rev 94(4):991-1026.

10. Padmanabhan K & Urban NN (2014) Disrupting information coding via block of 4-AP-sensitive potassium channels. Journal of neurophysiology 112(5): 1054-1066.

11. Edgerton JR & Jaeger D (2011) Dendritic sodium channels promote active decorrelation and reduce phase locking to parkinsonian input oscillations in model globus pallidus neurons. J Neurosci 31(30):10919-10936.

12. Coulter DA & Carlson GC (2007) Functional regulation of the dentate gyrus by GABA-mediated inhibition. Prog Brain Res 163:235-243.

13. Dieni CV, Nietz AK, Panichi R, Wadiche JI, & Overstreet-Wadiche L (2013) Distinct determinants of sparse activation during granule cell maturation. J Neurosci 33(49):19131-19142.

14. Marin-Burgin A, Mongiat LA, Pardi MB, & Schinder AF (2012) Unique Processing During a Period of High Excitation/Inhibition Balance in Adult-Born Neurons. Science 335(6073):1238-1242.

15. Wang S, Scott BW, & Wojtowicz JM (2000) Heterogenous properties of dentate granule neurons in the adult rat. J Neurobiol 42(2):248-257.

16. Severa W, Parekh O, James CD, & Aimone JB (2017) A Combinatorial Model for Dentate Gyrus Sparse Coding. Neural Comput 29(1):94-117.

17. Goard M & Dan Y (2009) Basal forebrain activation enhances cortical coding of natural scenes. Nat Neurosci 12(11): 1444-1449.

18. Lledo PM & Valley M (2016) Adult Olfactory Bulb Neurogenesis. Cold Spring Harb Perspect Biol 8(8).

19. Deng W, Aimone JB, & Gage FH (2010) New neurons and new memories: how does adult hippocampal neurogenesis affect learning and memory? Nature reviews. Neuroscience 11(5):339-350.

20. Kropff E, Yang SM, & Schinder AF (2015) Dynamic role of adult-born dentate granule cells in memory processing. Current opinion in neurobiology 35:21-26.

21. Yassa MA & Stark CE (2011) Pattern separation in the hippocampus. Trends Neurosci 34(10): 515-525.

22. Leutgeb JK, Leutgeb S, Moser MB, & Moser EI (2007) Pattern separation in the dentate gyrus and CA3 of the hippocampus. Science 315(5814):961-966.

23. Anderson P, Morris R, Amaral D, Bliss TV, & O’Keefe J (2007) The Hippocampus Book (Oxford University Press).

24. Imayoshi I, Sakamoto M, Ohtsuka T, Takao K, Miyakawa T, Yamaguchi M, Mori K, Ikeda T, Itohara S, & Kageyama R (2008) Roles of continuous neurogenesis in the structural and functional integrity of the adult forebrain. Nat Neurosci 11(10): 1153-1161.

25. Aimone JB, Wiles J, & Gage FH (2009) Computational influence of adult neurogenesis on memory encoding. Neuron 61(2):187-202.

26. Renno-Costa C, Lisman JE, & Verschure PF (2010) The mechanism of rate remapping in the dentate gyrus. Neuron 68(6): 1051-1058.

27. Marr D (1971) Simple memory: a theory for archicortex. Philos Trans R Soc Lond B Biol Sci 262(841):23-81.

28. Treves A & Rolls ET (1994) Computational analysis of the role of the hippocampus in memory. Hippocampus 4(3):374-391.

29. Foster WR, Ungar LH, & Schwaber JS (1993) Significance of conductances in Hodgkin-Huxley models. Journal of neurophysiology 70(6):2502-2518.

30. Goldman MS, Golowasch J, Marder E, & Abbott LF (2001) Global structure, robustness, and modulation of neuronal models. J Neurosci 21(14):5229-5238.

31. Rathour RK & Narayanan R (2014) Homeostasis of functional maps in active dendrites emerges in the absence of individual channelostasis. Proc Natl Acad Sci U S A 111(17):E1787-1796.

32. Prinz AA, Bucher D, & Marder E (2004) Similar network activity from disparate circuit parameters. Nat Neurosci 7(12): 1345-1352.

33. Aradi I & Holmes WR (1999) Role of multiple calcium and calcium-dependent conductances in regulation of hippocampal dentate granule cell excitability. J Comput Neurosci 6(3):215-235.

34. Krueppel R, Remy S, & Beck H (2011) Dendritic integration in hippocampal dentate granule cells. Neuron 71(3):512-528.

35. Lubke J, Frotscher M, & Spruston N (1998) Specialized electrophysiological properties of anatomically identified neurons in the hilar region of the rat fascia dentata. Journal of neurophysiology 79(3): 1518-1534.

36. Mott DD, Turner DA, Okazaki MM, & Lewis DV (1997) Interneurons of the dentate-hilus border of the rat dentate gyrus: morphological and electrophysiological heterogeneity. J Neurosci 17(11):3990-4005.

37. Santhakumar V, Aradi I, & Soltesz I (2005) Role of mossy fiber sprouting and mossy cell loss in hyperexcitability: a network model of the dentate gyrus incorporating cell types and axonal topography. Journal of neurophysiology 93(1):437-453.

38. Lee D, Lin BJ, & Lee AK (2012) Hippocampal place fields emerge upon single-cell manipulation of excitability during behavior. Science 337(6096):849-853.

39. Schmidt-Hieber C, Jonas P, & Bischofberger J (2004) Enhanced synaptic plasticity in newly generated granule cells of the adult hippocampus. Nature 429(6988): 184-187.

40. Liu S, Wang J, Zhu D, Fu Y, Lukowiak K, & Lu YM (2003) Generation of functional inhibitory neurons in the adult rat hippocampus. J Neurosci 23(3):732-736.

41. Rall W (1977) Core conductor theory and cable properties of neurons. Handbook of physiology. The nervous system. Cellular biology of neurons, ed Kandel ER (American physiological society, Bethesda, MD), Vol 1, pp 39-97.

42. Esposito MS, Piatti VC, Laplagne DA, Morgenstern NA, Ferrari CC, Pitossi FJ, & Schinder AF (2005) Neuronal differentiation in the adult hippocampus recapitulates embryonic development. J Neurosci 25(44):10074-10086.

43. Aimone JB & Gage FH (2011) Modeling new neuron function: a history of using computational neuroscience to study adult neurogenesis. The European journal of neuroscience 33(6): 1160-1169.

44. Dupret D, Fabre A, Dobrossy MD, Panatier A, Rodriguez JJ, Lamarque S, Lemaire V, Oliet SHR, Piazza PV, & Abrous DN (2007) Spatial learning depends on both the addition and removal of new hippocampal neurons. Plos Biology 5(8):1683-1694.

45. Tashiro A, Sandler VM, Toni N, Zhao CM, & Gage FH (2006) NMDA-receptor-mediated, cell-specific integration of new neurons in adult dentate gyrus. Nature 442(7105):929-933.

46. Alvarez DD, Giacomini D, Yang SM, Trinchero MF, Temprana SG, Büttner KA, Beltramone N, & Schinder AF (2016) A disynaptic feedback network activated by experience promotes the integration of new granule cells. Science 354(6311):459-465.

47. Aimone JB, Wiles J, & Gage FH (2006) Potential role for adult neurogenesis in the encoding of time in new memories. Nat Neurosci 9(6):723-727.

48. Edelman GM & Gally JA (2001) Degeneracy and complexity in biological systems. Proc Natl Acad Sci U S A 98(24):13763-13768.

49. Carnevale NT & Hines ML (2006) The NEURON Book (Cambridge University Press, Cambridge, UK).

50. Anirudhan A & Narayanan R (2015) Analogous synaptic plasticity profiles emerge from disparate channel combinations. J Neurosci 35(11):4691-4705.

51. Rathour RK & Narayanan R (2012) Inactivating ion channels augment robustness of subthreshold intrinsic response dynamics to parametric variability in hippocampal model neurons. J Physiol 590(Pt 22):5629-5652.

52. Schmidt-Hieber C, Jonas P, & Bischofberger J (2007) Subthreshold dendritic signal processing and coincidence detection in dentate gyrus granule cells. J Neurosci 27(31):8430-8441.

53. Chen C (2004) ZD7288 inhibits postsynaptic glutamate receptor-mediated responses at hippocampal perforant path-granule cell synapses. The European journal of neuroscience 19(3):643-649.

54. Magee JC (1998) Dendritic hyperpolarization-activated currents modify the integrative properties of hippocampal CA1 pyramidal neurons. J Neurosci 18(19):7613-7624.

55. Beck H, Ficker E, & Heinemann U (1992) Properties of two voltage-activated potassium currents in acutely isolated juvenile rat dentate gyrus granule cells. Journal of neurophysiology 68(6):2086-2099.

56. Ferrante M, Migliore M, & Ascoli GA (2009) Feed-forward inhibition as a buffer of the neuronal input-output relation. Proc Natl Acad Sci U S A 106(42):18004-18009.

57. Goldman DE (1943) Potential, Impedance, and Rectification in Membranes. The Journal of general physiology 27(1):37-60.

58. Hodgkin AL & Katz B (1949) The effect of sodium ions on the electrical activity of giant axon of the squid. J Physiol 108(1):37-77.

59. Poirazi P, Brannon T, & Mel BW (2003) Pyramidal neuron as two-layer neural network. Neuron 37(6):989-999.

60. Narayanan R & Johnston D (2010) The h current is a candidate mechanism for regulating the sliding modification threshold in a BCM-like synaptic learning rule. Journal of neurophysiology 104(2):1020-1033.

61. Destexhe A, Babloyantz A, & Sejnowski TJ (1993) Ionic mechanisms for intrinsic slow oscillations in thalamic relay neurons. Biophysical journal 65(4): 1538-1552.

62. Eliot LS & Johnston D (1994) Multiple components of calcium current in acutely dissociated dentate gyrus granule neurons. Journal of neurophysiology 72(2):762-777.

63. Boss BD, Peterson GM, & Cowan WM (1985) On the number of neurons in the dentate gyrus of the rat. Brain Res 338(1): 144-150.

64. Mishra P & Narayanan R (2015) High-conductance states and A-type K+ channels are potential regulators of the conductance-current balance triggered by HCN channels. Journal of neurophysiology 113(1):23-43.

65. Overstreet-Wadiche LS, Bromberg DA, Bensen AL, & Westbrook GL (2006) Seizures accelerate functional integration of adult-generated granule cells. J Neurosci 26(15):4095-4103.

66. Pedroni A, Minh do D, Mallamaci A, & Cherubini E (2014) Electrophysiological characterization of granule cells in the dentate gyrus immediately after birth. Front Cell Neurosci 8:44.

67. Johnston D & Wu SM (1995) Foundations of cellular neurophysiology (The MIT Press, Cambridge, Massachusetts).

68. Solstad T, Moser EI, & Einevoll GT (2006) From grid cells to place cells: a mathematical model. Hippocampus 16(12): 1026-1031.

